# deSAMBA: fast and accurate classification of metagenomics long reads with sparse approximate matches

**DOI:** 10.1101/736777

**Authors:** Gaoyang Li, Bo Liu, Yadong Wang

## Abstract

**Summary:** Long read sequencing technologies are promising to metagenomics studies. However, there is still lack of read classification tools to fast and accurately identify the taxonomies of noisy long reads, which is a bottleneck to the use of long read sequencing. Herein, we propose deSAMBA, a tailored long read classification approach that uses a novel sparse approximate match block (SAMB)-based pseudo alignment algorithm. Benchmarks on real datasets demonstrate that deSAMBA enables to simultaneously achieve fast speed and good classification yields, which outperforms state-of-the-art tools and has many potentials to cutting-edge metagenomics studies.

**Availability and Implementation:** https://github.com/hitbc/deSAMBA.

**Supplementary information:**

## 1 Introduction

Long read sequencing technologies are promising, as they enable to implement real-time and portable sequencing for metagenomics samples (Quick *et al.*, 2016). However, it is still non-trivial to implement fast and accurate classification for long reads to identify their taxonomies due to several technical issues such as serious sequencing errors, incomplete reference and common sequences in related genomes. As a fundamental step to characterize metagenomes, this is an important open problem and more advanced tools are on wide demand.

Most of state-of-the-art tools, such as Kraken (Wood and Salzberg, 2014), Centrifuge (Kim *et al.*, 2016), Kaiju (Menzel *et al.*, 2016), Meta-Othello (Liu *et al.*, 2018), are designed for short reads. Mainly, they use pseudo alignments, i.e., the exact or approximate matches from reads to reference as signals to achieve fast speed without loss of accuracy on short reads. However, they could fail at the high sequencing error rates (typically 10%-20%) of long reads. Long read aligners (Li, 2018) could be used as alternatives, however, they are still time-consuming which could be not well-suited to large scale datasets or real-time tasks.

Herein, we propose de Bruijn graph-based Sparse Approximate Match Block Analyzer (deSAMBA), a tailored long read classification approach using a novel sparse approximate match-based pseudo alignment algorithm. deSAMBA enables to simultaneously achieve good classification yields and fast speed, which is promising to breakthrough the bottleneck of metagenomics long read classification.

## 2 Methods

deSAMBA supports the reads produced by ONT and PacBio platforms. It uses Unitig-BWT data structure (Guan *et al.*, 2018) to index the unitigs of the de Bruijn graph of the reference sequences, and finds similar blocks between reads and reference through the index. These blocks are called as sparse approximate match blocks (SAMBs), as they are usually sparsely placed in reads and approximately matched to the reference. Mainly, deSAMBA classifies a read in four steps:

1. deSAMBA partitions the read into a series of segments, and for each of the segments, it finds the local region having most consecutive k-mer matches as a seed block;
2. for each of the seeding blocks, deSAMBA retrieves a set of maximal exact matches (MEMs) to the unitigs of the reference, and extends the MEMs to generate SAMBs by local alignment;
3. deSAMBA greedily merges the SAMBs, and further extends the merged SAMBs by sparse dynamic programming (SDP)-based pseudo alignment against local reference sequences;
4. deSAMBA scores the extended SAMBs and identifies the taxonomy entity of the read by the highest scored SAMB, meanwhile, it also outputs the SAMBs as the pseudo alignments of the reads.

Also refer to Supplementary Notes for more detailed information.

## 3 Results

We benchmarked deSAMBA, Centrifuge, Kaiju and Minimap2 with 145 real datasets from single genomes (86 ONT datasets and 59 PacBio datasets, Supplementary Table 1), using a pseudo metagenomics approach (Kim *et al.*, 2016). 16284 complete genomes (8621 bacterial, 251 archaea and 7412 viral genomes, totally ∼35 Gbp) from NCBI RefSeq database were used as reference. There are 49 ONT and 22 PacBio datasets whose ground truth genomes are in the reference (these datasets are called as “WR-datasets”), and for each of the other 37 ONT and 37 PacBio datasets, there is at least one genome in the reference that has common ancestry at species or genus level to its ground truth genome (these datasets are called as “NR-datasets”). The sensitivity, accuracy, F1-measure and speed of read classification were assessed (Fig. 1).

**Fig. 1.**
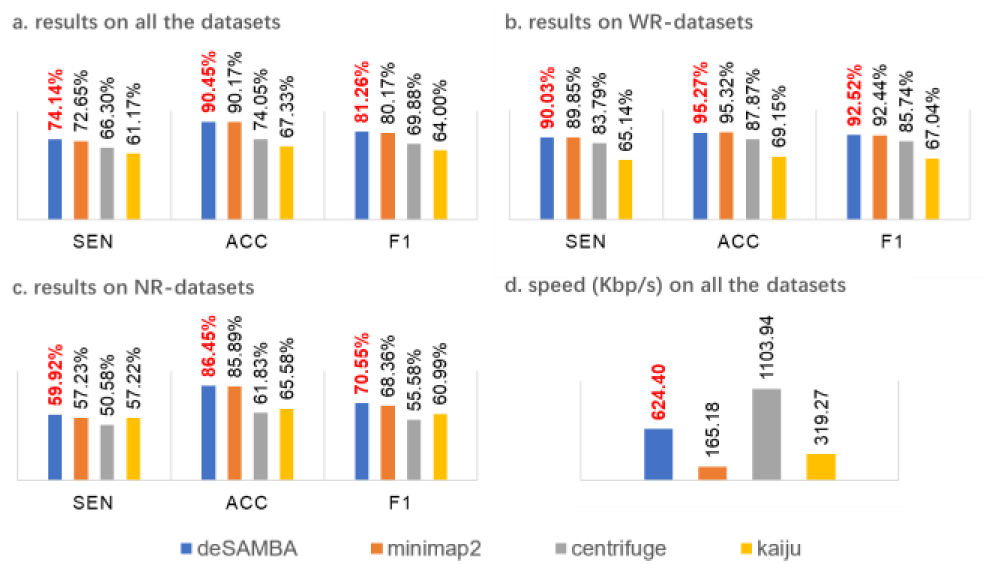
The results of the various approaches on the 145 real datasets. The first three subfigures respectively indicate the average sensitivity (“SEN”), accuracy (“ACC”) and F1-measure (“F1”) on all the 145 datasets (a), the 71 WR-datasets (b) and the 74 NR-datasets (c). The sensitivity, accuracy and F1-score are defined as S% = *N*_*TP*_*/N*_*T*_, A*% = N*_*TP*_*/N*_*R*_ and F1 = 2*SA*/(*S* + *A*) respectively, where *N*_*T*_, *N*_*R*_ and *N*_*TP*_ are respectively the total numbers of all the reads, the reads being classified and the reads being correctly classified. Herein, a read is considered as correctly classified only if its primary classification is assigned to its ground truth genome (if the read is from a WR-dataset), or a reference genome having the common ancestry at lowest level (species or genus, depending on the dataset) with its ground truth genome (if the read is from a NR-dataset). Hie speed of the approaches (d) is assessed by Kbp processed per second with 8 CPU threads.

Mainly, four issues are observed from the results.

Firstly, deSAMBA has good classification yields. deSAMBA has slightly better classification yields (F1-measure) than that of Minimap2, and both of them obviously outperformed the two pseudo-alignment-based approaches, Centrifuge and Kaiju, on both of sensitivity and accuracy (Fig. 1a and Supplementary Fig. 1a-b). Further, we assessed the classifications on the WR-datasets (Fig. 1b and Supplementary Fig. 1c-d), and similar trends were observed. These results indicate that, Centrifuge and Kaiju are easier to be affected by sequencing errors, even if the reads are from known genomes. This is mainly due to that such short read toward approaches rely on the hypothesis of long exact matches or few divergences between reads and reference, which does not stand for long reads. However, with the approximate matching approach, SAMBs can used as a kind of noise-robust features which enables deSAMBA find the evidence from noisy long reads to implement precise classification.

Secondly, deSAMBA has good ability to classify the reads from unknown genomes. We assessed the classifications on the NR-datasets (Fig. 1c and Supplementary Fig. 1e-f), and observed that deSAMBA has the best yields. We further investigated the intermediate results of deSAMBA, and found that the generated SAMBs usually have relatively large lengths and low edit distances to local reference sequences. Most of them can be specifically mapped to the reference genomes closely related to the ground truth genomes of the datasets, which greatly helps to produce correct classifications. It is also worthnoting that all the approaches have reduced sensitivities and accuracies on NR-datasets, mainly due to the large divergences between the ground truth genomes and their related genomes in the reference.

Thirdly, deSAMBA enables to identify the reads from various strains. Many studies ask to identify the strains of the sample. To assess this ability, we did an additional assessment with 7 ONT datasets having strain-level labels (Supplementary Fig. 2). In this assessment, a read is considered as correctly classified only if it is assigned to its ground truth strain. The results demonstrate that deSAMBA outperformed Minimap2 and Kaiju on all the 7 datasets. Moreover, Centrifuge only classified these reads to species level, seems that it does not support this function.

Fourthly, deSAMBA has fast speed. deSAMBA is about 4 and 2 times faster than Minimap2 and Kaiju, respectively, although slower than Centrifuge (Fig. 1d and Supplementary Fig. 1g-h). Considering its high sensitivity and accuracy, deSAMBA achieves a better balance between yields and speed than state-of-the-art pseudo alignment-based approaches, and it is more suited to large-scale datasets and real time tasks than long read aligners. Moreover, the memory footprint of deSAMBA is 69GB. Comparing to that of Centrifuge (9GB), Kaiju (31GB) and Minimap2 (76GB), this is acceptable, especially for modern servers.

## Supporting information

reference describe

