## Supplementary Material for "deSAMBA: fast and accurate classification of metagenomics long reads with sparse approximate matches"

Gaoyang Li[1, *], Bo Liu[1, *] and Yadong Wang[1, †]

[1.]Center for Bioinformatics, School of Computer Science and Technology, Harbin Institute of Technology, Harbin, Heilongjiang 150001, China

* The authors should be regarded as Joint First Authors.

† To whom correspondence should be addressed.

**Supplementary Table 1. The real datasets used for benchmark**

| # | SRA accession[a] | Sequencing Platform[b] | # of Reads[c] | # of Bases (Mbp)[d] | Organism[e] | Taxonomy ID[f] | Genus ID[g] | With Reference[h] |
|---|---|---|---|---|---|---|---|---|
| 1 | DRR129651 | ONT | 116592 | 744.10 | *Vibrio litoralis DSM 17657* | 1123493 | 662 | no |
| 2 | DRR129656 | ONT | 38252 | 464.56 | *Vibrio aphrogenes* | 1891186 | 662 | no |
| 3 | DRR129658 | ONT | 125964 | 910.34 | *Vibrio algivorus* | 1667024 | 662 | no |
| 4 | DRR129662 | ONT | 132936 | 801.39 | *Vibrio rumoiensis* | 76258 | 662 | no |
| 5 | DRR131199 | ONT | 44741 | 364.29 | *Arthrobacter sp. MN05-02* | 1571833 | 1663 | no |
| 6 | DRR154113 | ONT | 270603 | 1838.80 | *Streptococcus agalactiae* | 1311 | 1301 | yes |
| 7 | DRR154115 | ONT | 46024 | 195.03 | *Streptococcus pneumoniae* | 1313 | 1301 | yes |
| 8 | DRR164908 | ONT | 33965 | 233.14 | *Leptotrichia trevisanii* | 109328 | 32067 | no |
| 9 | DRR164910 | ONT | 62626 | 273.77 | *Leptotrichia wadei* | 157687 | 32067 | no |
| 10 | DRR164912 | ONT | 145853 | 598.02 | *Leptotrichia goodfellowii* | 157692 | 32067 | no |
| 11 | DRR164915 | ONT | 26286 | 126.45 | *Leptotrichia trevisanii* | 109328 | 32067 | no |
| 12 | ERR1358778 | ONT | 33642 | 285.16 | *Escherichia coli* | 562 | 561 | yes |
| 13 | ERR1418239 | ONT | 2547 | 10.57 | *Escherichia coli K-12* | 83333 | 561 | yes |
| 14 | ERR1769944 | ONT | 10102 | 166.54 | *GlucoNObacter oxydans 621H* | 290633 | 441 | yes |
| 15 | ERR2109178 | ONT | 7908 | 114.92 | *Enterococcus faecium* | 1352 | 1350 | yes |
| 16 | ERR2195906 | ONT | 10039 | 27.90 | *Citrobacter koseri* | 545 | 544 | yes |
| 17 | ERR2259087 | ONT | 1533 | 25.10 | *Neisseria meningitidis* | 487 | 482 | yes |
| 18 | ERR2278795 | ONT | 104 | 0.68 | *Streptomyces cinnamoneus* | 53446 | 1883 | no |
| 19 | ERR2722109 | ONT | 288320 | 1919.33 | *Campylobacter jejuni subsp. Jejuni* | 32022 | 194 | yes |
| 20 | ERR2724040 | ONT | 69052 | 166.18 | *Mycobacterium tuberculosis* | 1773 | 1763 | yes |
| 21 | ERR776852 | ONT | 2312 | 18.77 | *Acinetobacter sp. ADP1* | 62977 | 469 | no |
| 22 | ERR977574 | ONT | 7202 | 48.23 | *Bacteroides fragilis* | 817 | 816 | yes |
| 23 | SRR3191595 | ONT | 29682 | 170.98 | *Agrobacterium tumefaciens* | 358 | 357 | yes |
| 24 | SRR4046732 | ONT | 4346 | 22.20 | *Xanthomonas citri pv. malvacearum* | 86040 | 338 | yes |
| 25 | SRR5117441 | ONT | 27892 | 217.97 | *Yersinia pestis* | 632 | 629 | yes |
| 26 | SRR5344353 | ONT | 2680 | 14.84 | *Akkermansia muciniphila* | 239935 | 239934 | yes |
| 27 | SRR5344354 | ONT | 21368 | 50.89 | *LachNOclostridium sp. An76* | 1965654 | 1506553 | no |
| 28 | SRR5344355 | ONT | 17954 | 44.43 | *Anaerotignum lactatifermentans* | 160404 | 2039240 | no |

| | | | | | | | | |
|---|---|---|---|---|---|---|---|---|
| 29 | SRR5344357 | ONT | 5455 | 34.66 | *Bacteroides sp. An51A* | 1965640 | 816 | no |
| 30 | SRR5344358 | ONT | 15973 | 56.64 | *Flavonifractor sp. An10* | 1965537 | 946234 | no |
| 31 | SRR5429682 | ONT | 9174 | 68.80 | *Klebsiella pneumoniae subsp. Pneumoniae* | 72407 | 570 | yes |
| 32 | SRR5457530 | ONT | 6139 | 36.21 | *Clostridioides difficile* | 1496 | 1870884 | yes |
| 33 | SRR5514549 | ONT | 12219 | 72.91 | *Achromobacter denitrificans* | 32002 | 222 | yes |
| 34 | SRR5629775 | ONT | 68851 | 808.43 | *Klebsiella oxytoca* | 571 | 570 | yes |
| 35 | SRR5629779 | ONT | 6948 | 89.05 | *Serratia marcescens* | 615 | 613 | yes |
| 36 | SRR5665595 | ONT | 61681 | 836.52 | *Klebsiella pneumoniae* | 573 | 570 | yes |
| 37 | SRR5891470 | ONT | 19378 | 237.24 | *Acinetobacter baumannii* | 470 | 469 | yes |
| 38 | SRR5997379 | ONT | 9345 | 33.01 | *Mesoplasma chauliocola* | 216427 | 46239 | yes |
| 39 | SRR6082028 | ONT | 23223 | 228.70 | *Mesoplasma lactucae ATCC 49193* | 81460 | 46239 | yes |
| 40 | SRR6129218 | ONT | 93588 | 783.29 | *Mesoplasma florum* | 2151 | 46239 | yes |
| 41 | SRR6224382 | ONT | 21302 | 177.54 | *Mesoplasma entomophilum* | 2149 | 46239 | yes |
| 42 | SRR6312194 | ONT | 23297 | 49.53 | *Paenibacillus pasadenensis* | 217090 | 44249 | no |
| 43 | SRR6327830 | ONT | 25654 | 155.58 | *Legionella sainthelensi* | 28087 | 445 | yes |
| 44 | SRR6364637 | ONT | 133722 | 363.43 | *Mesoplasma syrphidae* | 225999 | 46239 | yes |
| 45 | SRR6471048 | ONT | 20039 | 383.52 | *Mycoplasma hominis* | 2098 | 2093 | yes |
| 46 | SRR6475285 | ONT | 45031 | 644.93 | *Streptomyces lunaelactis* | 1535768 | 1883 | no |
| 47 | SRR6780924 | ONT | 33201 | 165.87 | *Fusobacterium periodonticum* | 860 | 848 | yes |
| 48 | SRR6830111 | ONT | 17411 | 105.82 | *Fusobacterium nucleatum subsp. nucleatum* | 76856 | 848 | yes |
| 49 | SRR6917534 | ONT | 18610 | 121.69 | *Streptococcus pyogenes* | 1314 | 1301 | yes |
| 50 | SRR7119552 | ONT | 55045 | 332.69 | *Acinetobacter NOsocomialis* | 106654 | 469 | yes |
| 51 | SRR7119560 | ONT | 42530 | 308.59 | *Acinetobacter pittii* | 48296 | 469 | yes |
| 52 | SRR7467298 | ONT | 265157 | 2370.25 | *AmiNObacter sp. MSH1* | 374606 | 31988 | no |
| 53 | SRR7467447 | ONT | 445670 | 2554.30 | *Streptomyces sp.* | 1931 | 1883 | no |
| 54 | SRR7532470 | ONT | 58457 | 664.97 | *Clostridium botulinum* | 1491 | 1485 | yes |
| 55 | SRR7739756 | ONT | 70760 | 685.10 | *Staphylococcus aureus* | 1280 | 1279 | yes |
| 56 | SRR7908033 | ONT | 46226 | 1292.88 | *Bordetella pertussis* | 520 | 517 | yes |
| 57 | SRR7957428 | ONT | 364969 | 968.02 | *CapNOcytophaga canimorsus* | 28188 | 1016 | yes |
| 58 | SRR7989235 | ONT | 108976 | 839.33 | *Borrelia miyamotoi* | 47466 | 138 | yes |

| | | | | | | | | |
|---|---|---|---|---|---|---|---|---|
| 59 | SRR8030961 | ONT | 25668 | 171.13 | *Microbacterium foliorum* | 104336 | 33882 | no |
| 60 | SRR8069226 | ONT | 275794 | 2176.62 | *Thalassotalea euphylliae* | 1655234 | 1518149 | no |
| 61 | SRR8081954 | ONT | 9451 | 76.11 | *SiNOrhizobium meliloti* | 382 | 28105 | yes |
| 62 | SRR8112132 | ONT | 201287 | 2248.28 | *Achromobacter sp. B7* | 2282475 | 222 | no |
| 63 | SRR8113455 | ONT | 2,174,450 | 11400.00 | *Thermoanaerobacter ethaNOlicus JW 200* | 509192 | 1754 | no |
| 64 | SRR8115246 | ONT | 2,404,454 | 6100.00 | *Pantoea agglomerans* | 549 | 53335 | yes |
| 65 | SRR8185373 | ONT | 39767 | 545.95 | *Rhodococcus sp. P1Y* | 1302308 | 1827 | no |
| 66 | SRR8278838 | ONT | 110157 | 268.48 | *Helicobacter pylori* | 210 | 209 | yes |
| 67 | SRR8306005 | ONT | 75727 | 857.01 | *Wolbachia endosymbiont of Drosophila ananassae* | 307502 | 953 | no |
| 68 | SRR8335319 | ONT | 84212 | 1092.09 | *Vibrio campbellii DS40M4* | 1088888 | 662 | yes |
| 69 | SRR8362629 | ONT | 168429 | 767.51 | *Mannheimia varigena* | 85404 | 75984 | no |
| 70 | SRR8428672 | ONT | 159650 | 1122.23 | *Ochrobactrum intermedium* | 94625 | 528 | no |
| 71 | SRR8457080 | ONT | 8417 | 43.56 | *Neisseria goNOrrhoeae* | 485 | 482 | yes |
| 72 | SRR8467877 | ONT | 525599 | 2633.69 | *Caldicellulosiruptor changbaiensis* | 1222016 | 44000 | no |
| 73 | SRR8480439 | ONT | 69399 | 918.04 | *Brevundimonas diminuta* | 293 | 41275 | no |
| 74 | SRR8480530 | ONT | 234432 | 2365.95 | *Bacillus subtilis subsp. spizizenii ATCC 6633* | 96241 | 1386 | yes |
| 75 | SRR8494918 | ONT | 100105 | 1099.82 | *Citrobacter gillenii* | 67828 | 544 | no |
| 76 | SRR8536147 | ONT | 1,000,846 | 8300.00 | *Halomonas olivaria* | 390919 | 2745 | no |
| 77 | SRR8538950 | ONT | 64431 | 173.74 | *Staphylococcus pseudintermedius* | 283734 | 1279 | yes |
| 78 | SRR8549412 | ONT | 74931 | 429.56 | *Halomonas sulfidaeris* | 115553 | 2745 | no |
| 79 | SRR8550983 | ONT | 81829 | 749.91 | *Psychrobacter sp. KH172YL61* | 2517899 | 497 | no |
| 80 | SRR8554089 | ONT | 76024 | 510.07 | *Halomonas axialensis* | 115555 | 2745 | no |
| 81 | SRR8560598 | ONT | 11084 | 165.31 | *Sphingomonas paucimobilis* | 13689 | 13687 | no |
| 82 | SRR8664470 | ONT | 543832 | 4410.84 | *Klebsiella sp. PO552* | 1972757 | 570 | no |
| 83 | SRR8697008 | ONT | 19085 | 303.34 | *Brevundimonas naejangsanensis* | 588932 | 41275 | yes |
| 84 | SRR8749593 | ONT | 13605 | 25.91 | *Burkholderia pseudomallei* | 28450 | 32008 | yes |
| 85 | SRR8767485 | ONT | 2029 | 28.51 | *Enterobacter hormaechei* | 158836 | 547 | no |
| 86 | SRR8767486 | ONT | 6602 | 100.54 | *Citrobacter freundii* | 546 | 544 | yes |
| 87 | ERR1140956 | PacBio | 64922 | 823.64 | *Staphylococcus aureus* | 1280 | 1279 | yes |

| | | | | | | | | |
|---|---|---|---|---|---|---|---|---|
| 88 | ERR1354168 | PacBio | 100569 | 1257.47 | *Streptococcus sobrinus* | 1310 | 1301 | no |
| 89 | ERR1543212 | PacBio | 83955 | 1111.02 | *Acinetobacter junii* | 40215 | 469 | yes |
| 90 | ERR1599942 | PacBio | 11808 | 87.44 | *Campylobacter upsaliensis* | 28080 | 194 | no |
| 91 | ERR1802424 | PacBio | 88061 | 914.95 | *Actinomyces slackii* | 52774 | 1654 | no |
| 92 | ERR1805700 | PacBio | 65521 | 747.50 | *Mycoplasma hyorhinis* | 2100 | 2093 | yes |
| 93 | ERR1857497 | PacBio | 35380 | 396.41 | *Bartonella grahamii* | 33045 | 773 | no |
| 94 | ERR1883080 | PacBio | 115549 | 1013.75 | *Helicobacter mustelae* | 217 | 209 | no |
| 95 | ERR1940980 | PacBio | 82137 | 950.67 | *Helicobacter pullorum* | 35818 | 209 | no |
| 96 | ERR1940990 | PacBio | 95528 | 912.78 | *Clostridium paraputrificum* | 29363 | 1485 | no |
| 97 | ERR2125654 | PacBio | 84110 | 859.20 | *Kocuria rosea* | 1275 | 57493 | no |
| 98 | ERR2125846 | PacBio | 86844 | 1282.82 | *Achromobacter xylosoxidans* | 85698 | 222 | yes |
| 99 | ERR2146854 | PacBio | 110931 | 1257.07 | *Citrobacter youngae* | 133448 | 544 | no |
| 100 | ERR2146855 | PacBio | 89496 | 827.10 | *Actinobacillus equuli* | 718 | 713 | no |
| 101 | ERR2225503 | PacBio | 46696 | 551.10 | *Helicobacter fennelliae* | 215 | 209 | no |
| 102 | ERR2237784 | PacBio | 16373 | 123.42 | *Campylobacter ureolyticus* | 827 | 194 | no |
| 103 | ERR2237840 | PacBio | 115641 | 1343.08 | *Helicobacter cinaedi* | 213 | 209 | no |
| 104 | ERR2246641 | PacBio | 82270 | 782.65 | *Actinomyces howellii* | 52771 | 1654 | no |
| 105 | ERR2246671 | PacBio | 119321 | 1355.07 | *Shigella boydii* | 621 | 620 | no |
| 106 | ERR2432878 | PacBio | 56894 | 561.80 | *Acinetobacter calcoaceticus* | 471 | 469 | yes |
| 107 | ERR2532027 | PacBio | 79831 | 870.43 | *Neisseria gonorrhoeae* | 485 | 482 | yes |
| 108 | ERR2532043 | PacBio | 71229 | 518.42 | *Listeria innocua* | 1642 | 1637 | no |
| 109 | ERR2532096 | PacBio | 33466 | 299.86 | *Clostridium tertium* | 1559 | 1485 | no |
| 110 | ERR2532135 | PacBio | 121764 | 1175.98 | *Vibrio furnissii* | 29494 | 662 | no |
| 111 | ERR2532412 | PacBio | 84519 | 862.70 | *Actinobacillus pleuropneumoniae* | 715 | 713 | no |
| 112 | ERR2543055 | PacBio | 126847 | 1260.59 | *Shewanella putrefaciens* | 24 | 22 | no |
| 113 | ERR2749075 | PacBio | 102245 | 980.09 | *Mycobacteroides abscessus* | 36809 | 1763 | yes |
| 114 | ERR772459 | PacBio | 81190 | 894.16 | *Citrobacter koseri* | 545 | 544 | yes |
| 115 | ERR772462 | PacBio | 139046 | 922.92 | *Actinobacillus ureae* | 723 | 713 | no |
| 116 | ERR841690 | PacBio | 81618 | 1040.88 | *Neisseria lactamica* | 486 | 482 | yes |
| 117 | ERR910531 | PacBio | 152267 | 1110.66 | *Streptococcus sanguinis* | 1305 | 1301 | no |
| 118 | SRR2163073 | PacBio | 126276 | 1203.68 | *Bacillus licheniformis* | 1402 | 1386 | yes |
| 119 | SRR2560502 | PacBio | 57657 | 140.32 | *Bacteroides vulgatus* | 821 | 816 | no |

| | | | | | | | | |
|---|---|---|---|---|---|---|---|---|
| 120 | SRR2925022 | PacBio | 117636 | 822.15 | *Clostridium botulinum* | 1491 | 1485 | yes |
| 121 | SRR4176232 | PacBio | 177858 | 704.60 | *Carnobacterium iners* | 1073423 | 2747 | no |
| 122 | SRR4180946 | PacBio | 136430 | 608.67 | *Pseudarthrobacter chlorophenolicus* | 85085 | 1742993 | no |
| 123 | SRR4181438 | PacBio | 29049 | 131.32 | *Rhodococcus percolatus* | 45461 | 1827 | no |
| 124 | SRR4181588 | PacBio | 236611 | 802.53 | *Amycolatopsis niigatensis* | 369932 | 1813 | no |
| 125 | SRR4236930 | PacBio | 111645 | 474.70 | *Achromobacter sp. MFA1 R4* | 1881016 | 222 | no |
| 126 | SRR4237132 | PacBio | 231416 | 869.52 | *Terriglobus roseus* | 392734 | 392733 | no |
| 127 | SRR4244870 | PacBio | 118334 | 493.82 | *Bradyrhizobium erythrophlei* | 1437360 | 374 | no |
| 128 | SRR5165186 | PacBio | 43375 | 88.08 | *Mesorhizobium loti* | 381 | 68287 | no |
| 129 | SRR5165459 | PacBio | 167285 | 630.72 | *Mesorhizobium australicum* | 536018 | 68287 | no |
| 130 | SRR5320243 | PacBio | 44474 | 146.97 | *Mycoplasma hyopneumoniae* | 2099 | 2093 | yes |
| 131 | SRR5829900 | PacBio | 145413 | 619.16 | *Clostridium beijerinckii* | 1520 | 1485 | yes |
| 132 | SRR6256365 | PacBio | 104731 | 462.71 | *Caldicellulosiruptor bescii* | 31899 | 44000 | no |
| 133 | SRR7072821 | PacBio | 160498 | 640.46 | *Halanaerobium congolense* | 54121 | 2330 | no |
| 134 | SRR7135411 | PacBio | 69488 | 378.16 | *Acidovorax citrulli* | 80869 | 12916 | yes |
| 135 | SRR7637788 | PacBio | 105804 | 401.74 | *Helicobacter saguini* | 1548018 | 209 | no |
| 136 | SRR7967879 | PacBio | 92278 | 1108.33 | *Borreliella burgdorferi* | 139 | 64895 | yes |
| 137 | SRR8175749 | PacBio | 65808 | 1006.98 | *Acinetobacter haemolyticus* | 29430 | 469 | yes |
| 138 | SRR8270580 | PacBio | 254965 | 597.83 | *Xanthomonas vasicola* | 56459 | 338 | no |
| 139 | SRR8281303 | PacBio | 119161 | 844.92 | *Deinococcus wulumuqiensis* | 980427 | 1298 | no |
| 140 | SRR8292561 | PacBio | 93819 | 1382.05 | *Salmonella enterica* | 28901 | 590 | yes |
| 141 | SRR8293937 | PacBio | 215678 | 1066.08 | *Streptococcus agalactiae* | 1311 | 1301 | yes |
| 142 | SRR8436601 | PacBio | 31434 | 66.13 | *Escherichia coli* | 562 | 561 | yes |
| 143 | SRR8476224 | PacBio | 65455 | 416.61 | *Bacillus coagulans* | 1398 | 1386 | yes |
| 144 | SRR8494910 | PacBio | 145862 | 1239.14 | *Citrobacter braakii* | 57706 | 544 | yes |
| 145 | SRR8691672 | PacBio | 48351 | 441.98 | *Acinetobacter baumannii* | 470 | 469 | yes |

a) The SRA accession number of the dataset;

b) The type of sequencing platform produces the dataset (ONT or PacBio);

c) The number of reads involved in the dataset;

d) The total number of bases of the reads in the dataset, counted by million basepairs (Mbp);

e) The organism of the sample of the dataset, i.e., the ground truth genome the dataset being from;

f) The taxonomy ID of the organism of the dataset;

g) The genus ID of the organism of the dataset;

h) Whether the ground truth genome sequence is in the reference (yes or no).

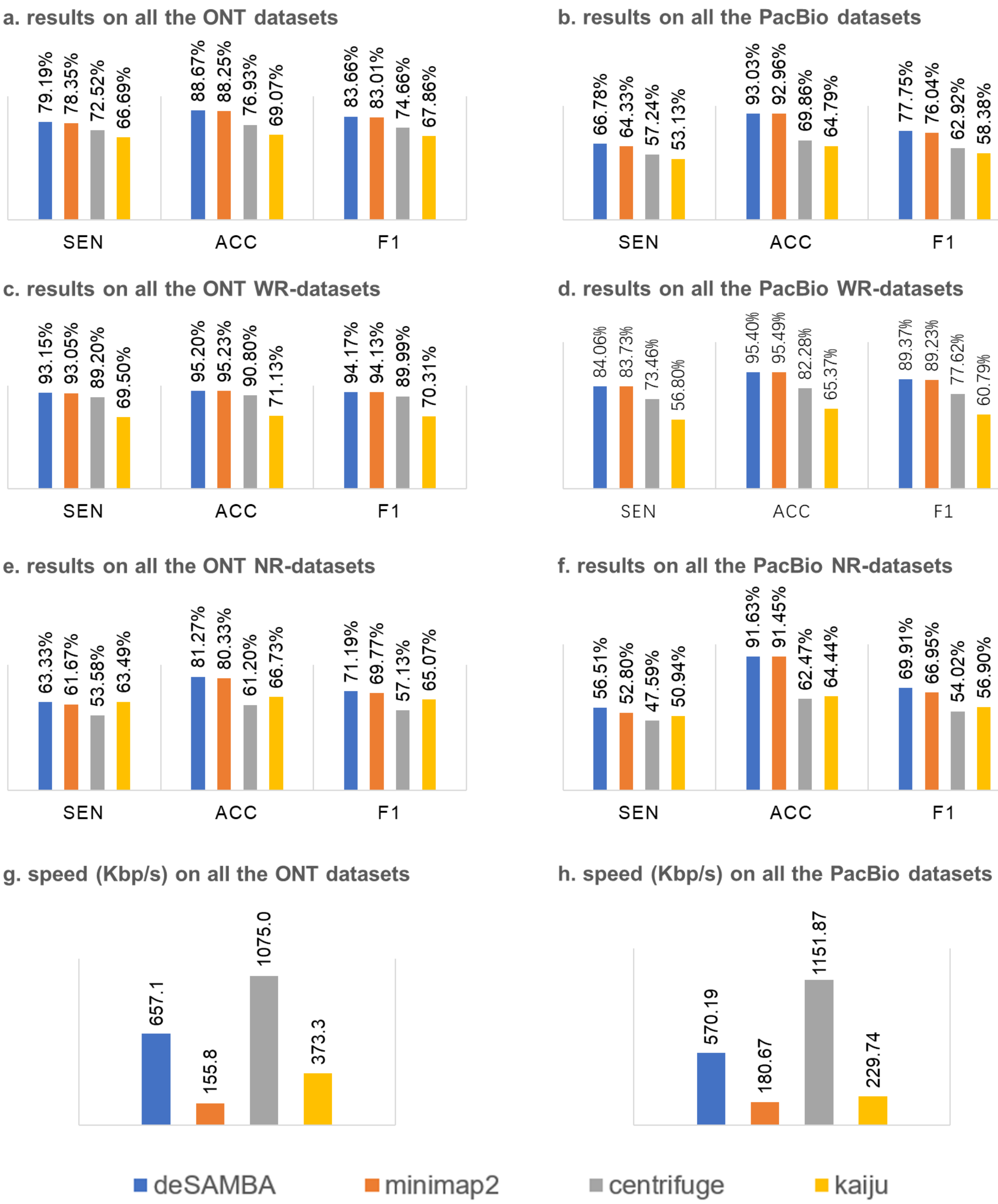

**Supplementary Fig 1. The results of the various approaches on the 86 ONT and 59 PacBio datasets**

The subfigures a-f respectively indicate the average sensitivity ("SEN"), accuracy ("ACC"), F1-measure ("F1") on all the ONT (a) and PacBio (b) datasets, the WR-datasets produced by ONT (c) and PacBio (d) platforms, the NR-datasets produced by ONT (e) and PacBio (f) platforms. The speed of the various approaches on all the 86 ONT and 59 PacBio datasets are respectively shown in subfigures g and h (assessed with 8 CPU threads).

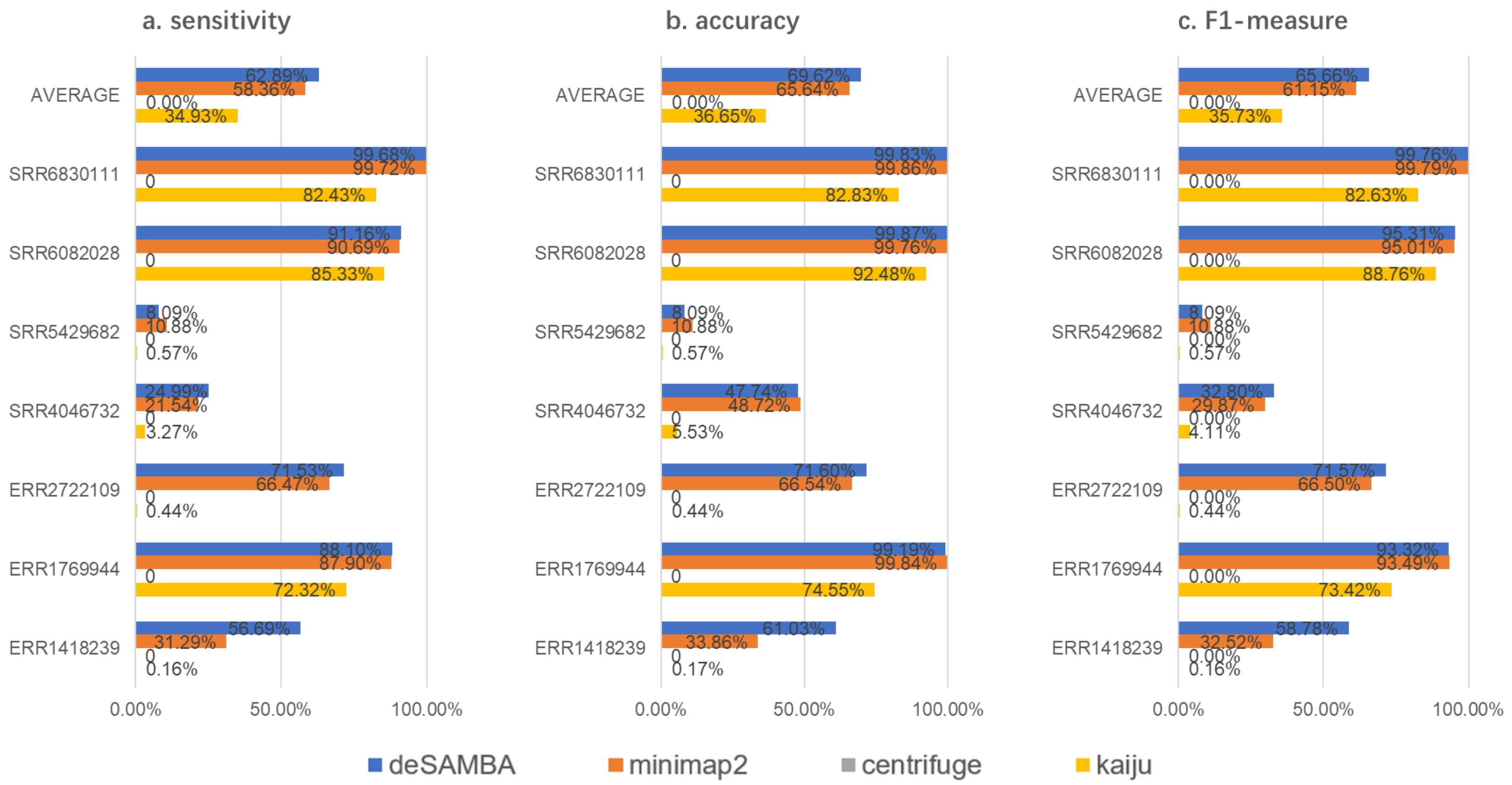

**Supplementary Fig 2. The results of the various approaches on the 7 ONT datasets with strain-level labels**

The sensitivity (a), accuracy (b) and F1-Measure (c) of the four approaches on the 7 ONT datasets with strain-level labels. The averaged statistics (marked as "AVERAGE") and the statistics on the 7 datasets (SRA Accessions: ERR1418239, ERR1769944, ERR2722109, SRR4046732, SRR5429682, SRR6082028 and SRR6830111, corresponding to lines 13, 14, 19, 24, 31, 39, 48 of Supplementary Table 1) are respectively shown in the bar plots.

### Supplementary Notes

#### 1. Supplementary Methods

The classification of metagenomics long reads is a non-trivial task due to several technical issues: 1) most of the long reads produced by mainstream platforms (such as ONT and PacBio platforms) are error-prone, which requires the read classifier to be noise-robust; 2) the reference is usually incomplete, i.e., many reads could be from unknown genomes, which requires the read classifier to be able to handle the divergences between the ground truth genomes and their related genomes in the reference; 3) there are usually many common sequences among the closely related genomes (e.g., the genomes of the various strains of a bacteria species) in the reference, which requires the read classifier to be able to handle the large amount of repeats in the reference. Under such circumstance, exact match-based pseudo alignment approaches could not work well, since there would be very few long exact matches between the read and the reference due to the sequencing errors, and short exact matches have poor ability to handle the ubiquitous repeats in the reference. Approximate match-based pseudo alignment could be a feasible way, since it enables to be tolerant to the divergences between the read and the reference, however, tailored design and implementation are still needed to handle many practical issues in metagenomics long read datasets.

deSAMBA is an approximate match-based pseudo alignment approach specifically designed for long read classification. It is motivated by the results of the previous study (Chaisson and Tesler, 2012) that sequencing errors are unevenly distributed along the reads. Many long-read aligners (Chaisson and Tesler, 2012; Li, 2013; Li, 2018; Sedlazeck *et al.*, 2018) also take advantage of this model to find the exact matches between read and reference to compose seeding-and-extension approaches. However, such seeds have limited lengths which could be not suited to large and repetitive reference sequences. Other than exact matches, the main idea of deSAMBA is to find the read blocks having relatively large lengths and low edit distances to local reference sequences. Previous studies indicate that such long approximated matching blocks usually can be specifically mapped to few genomics positions with high accuracy (Liu *et al.*, 2017), so that we realize they can be used as a kind of noise-robust feature to implement precise read classification.

deSAMBA is composed by a series of tailored designs and implementations to simultaneously achieve fast speed and excellent classification yields. Some detailed information is as following.

##### 1.1 The indexing of reference sequences

deSAMBA organizes the reference sequences in a de Bruijn graph-based approach which is initially proposed in deBGA (Liu *et al*., 2016). To reduce memory footprint, we used Unitig-BWT data structure (Guan *et al*., 2018) to index the unitig of the de Bruijn graph of the given reference sequences. More precisely, deSAMBA constructs the de Bruijn graph of reference sequences at first, and extracts the unitigs. The Burrows-Wheeler Transformation is then constructed for the concatenated unitigs as the Unitig-BWT of the reference sequences. This indexing approach has good balance between retrieval speed and RAM space cost. It is also worthnoting that, instead of assigning a unique taxonomy ID like the previous study (Liu *et al*., 2016), deSAMBA maintains a position list for each of the unitigs. For a certain unitig, each item of the position list records a genomic position in reference, which represents the location of a specific copy of the unitig.

In addition to the UNITIG-BWT index, deSAMBA also builds a bloom filter-based index, which is used as an auxiliary index to the generation of SAMBs. The bloom filter-based index enables to give a quick answer whether a k-mer in a given read appears in the reference, and it helps to fast find the candidate positions which are likely to be within the read blocks highly similar to local reference sequences.

With the Unitig-BWT index and the auxiliary bloom filter-based index, deSAMBA classifies a given read in the following four steps.

##### 1.2 The generation of seed blocks

deSAMBA initially partitions a given read into 100 bp long segments. For each of the segments, deSAMBA exacts all its k-mers, and separately test each of the k-mer with the bloom filter-based index. The k-mers passed the bloom filter are recorded, and deSAMBA finds the local region in the segment having most consecutive passed k-mers, i.e., the longest passed k-mer chain, as a “seed block”. It is also worthnoting that the k parameter is automatically configured in advance according to the size of the reference, and it is set as 15-19 bp in most of the cases.

This idea derives from the characteristics of bloom filter. With a bloom filter-based index, there could be a proportion of false positives, i.e., a passed k-mer could be a false positive “hit” to reference. However, the false positive rate is relatively low, so that a seed block is more likely to be a true positive >kbp long exact match to the reference than false positive k-mers. Thus, deSAMBA assumes that the seed blocks are long exact matches, and uses them as candidates to generate SAMBs.

Moreover, it is not problematic if a seed block is a false positive match, since deSAMBA would retrieve the corresponding sequence of the seed block through the Unitig-BWT index and filter it out if no such sequence can be found.

**1.3 The generation of initial SAMBs**

For each of the seed blocks, deSAMBA exacts all the suffixes of the seed block, and efficiently retrieves the maximal exact matches between the suffixes and the unitigs of the reference through the Unitig-BWT index. All the retrieved matches are called as U-MEMs, and deSAMBA separately checks each of them. If a U-MEM can be fully covered by another one, deSAMBA would filter it out. After the filtration, the longest eight remining U-MEMs are used to generate initial SMABs.

For each of the U-MEMs, deSAMBA maps it to all the copies of its matched unitig, i.e., the U-MEM are converted to one or more MEMs to local reference sequences (each of them is called as a "R-MEM"). And for each of the R-MEMs, deSAMBA uses Landau-Vishkin algorithm to compose an alignment between the read part and the local reference to extend the R-MEM to a longer approximate match block. The extension is limited to the flanking 12bp of the local reference sequence of the R-MEM, and it is expected that the alignment has a low edit distance, and a quality score is assigned to the generated approximate match block. The quality score is calculated based on all the matched and mismatched bases in the alignment with the following equations.

$$S = \sum_{\substack{matched \\ bases}} S_{match} \times N_{match} + \sum_{\substack{mismatched \\ bases}} S_{mis} \times N_{mis} + S_{Ref\_Complex\_Penalty}$$

$$S_{match} = -10 \times \lg(\frac{0.25}{1 - \mathrm{E}})$$

$$S_{mis} = -10 \times \lg(\frac{0.75}{\mathrm{E}})$$

$$S_{Ref\_Complex\_Penalty} = -10 \times \lg(\mathrm{L}_{reference})$$

Herein, $S_{match}$ and $S_{mis}$ are the scores of matched and mismatched bases, $S_{Ref\_Complex\_Penalty}$ is a reference size-based penalty which is related to the total length of the reference $\mathrm{L}_{reference}$, $N_{match}$ and $N_{mis}$ are respectively the numbers of matched bases and mismatched bases in the alignment, and *E* is a parameter representing the expected sequencing error rate (default value: 0.15).

After scoring, the generated approximate match blocks having >30 quality scores are remained as "initial SAMBs", and other ones are discarded. Moreover, each of the SAMBs can be written as a 4-tuple: $SAMB_i = (r_i^S, r_i^E, R_i^S, R_i^E)$, where $r_i^S$ and $r_i^E$ are the start and end positions on the read, and $R_i^S$

and $R_i^E$ are the start and end positions on the reference, respectively.

**1.4 The generation of extended SAMBs**

deSAMBA greedily merges initial SAMBs from upstream to downstream. Two SAMBs are combined if they are distanced less than 300 bp on the reference, and the difference between their distances on the read and on the reference is less than 30 bp. After this processing, the initial SAMBs are combined as a series of SAMB-chains. Each SAMB-chain can be written as a series of SAMBs:

$$SC_i = \{SAMB_{ij}, j = 1 \dots |SC_i|\}$$

where $|SC_i|$ is the number of SAMBs of $SC_i$. Moreover, it can be derived that the read part and the local reference sequence covered by a SAMB-chain $SC_i$ are $[r_{i1}^S, r_{i|SC_i|}^E]$ and $[R_{i1}^S, R_{i|SC_i|}^E]$, respectively.

deSAMBA further extends the SAMB-chains by a sparse dynamic programming (SDP)-based pseudo alignment approach. For a SAMB-chain, $SC_i$, this is done in the following four sub-steps.

1) deSAMBA extracts all the 9-mers within $[r_{i1}^S - LR, r_{i|SC_i|}^E + LR]$, and indexes them with a hash table-based data structure. The parameter *LR* (default value: 1000) defines an extended local region in reference.

2) deSAMBA retrieves all the 9bp matches between$[R_{i1}^S, R_{i|SC_i|}^E]$ and $[r_{i1}^S - LR, r_{i|SC_i|}^E + LR]$ through the 9-mer hash table, and combines all the consecutive 9-mer matches into one or more longer exact matches.

3) The remaining matches are chained in an SDP approach with the following function:

$$f(\mathrm{Mat}_p) = \max\{\max_{p>q\geq 1}\{f(\mathrm{Mat}_q) + L(\mathrm{Mat}_p) - 8 - \theta(p,q)\}, L(\mathrm{Mat}_p)\}$$

$$\theta(p,q) = \begin{cases} 0.1 \times \left((Mat_p^R - Mat_q^R) - (Mat_p^r - Mat_q^r)\right) & if\ Mat_p^R - Mat_q^R < 600 \\ \infty & otherwise \end{cases}$$

where $\mathrm{Mat}_p$ and $\mathrm{Mat}_q$ are the *p*-th and q-th matches (sorted by reference position) and they are not overlapped, $Mat_p^R$ and $Mat_q^R$ are their positions on the reference, $Mat_p^r$ and $Mat_q^r$ are their positions on the read respectively; $f(\mathrm{Mat}_p)$ is the scoring function for the $\mathrm{Mat}_p$, $L(\mathrm{Mat}_p)$ is the length of $\mathrm{Mat}_p$, and $\theta(p,q)$ is a penalty for the two linked matches, $\mathrm{Mat}_p$ and $\mathrm{Mat}_q$.

4) After the SDP, the optimal chain of matches is obtained through backtracking, and it recorded as the “extended SAMB” generated based on $SC_i$.

**1.5 The classification of the reads**

deSAMBA collects all the extended SAMBs generated, and sorts them by their scores calculated in the SDP process. deSAMBA then determines the primary classification of the read by the taxonomy entity of the reference genome corresponding to the extended SAMB with highest score. The taxonomy entities of other extended SAMBs are as secondary classifications. Moreover, the SAMBs are also output as the partial pseudo alignments of the read.

#### 2. Implementation of benchmark

All the benchmarks were carried out on a server with 4 Intel E7-4820 CPUs (32 cores) and 1 TB RAM running Ubuntu Linux OS. All the benchmarked classification tools were run in 8 CPU threads. Some detailed information about employed reference sequences, the real sequencing datasets and the command lines used for read classification is as following.

##### 2.1 Reference sequences used for benchmark

We downloaded all reference sequences from NCBI RefSeq database. A genome sequence from RefSeq database was employ only if it is marked as a “complete genome”. There are totally 8621 bacterial, 251 archaea and 7412 viral genomes being used. The RefSeq ID and Taxonomy ID are described in “reference describe.txt”. For kaiju, the reference index was built using NCBI protein database, due to its specifically designed read classification approach.

Downloading “assembly_summary” files:

ftp://ftp.ncbi.nlm.nih.gov/genomes/refseq/bacteria/assembly_summary.txt

ftp://ftp.ncbi.nlm.nih.gov/genomes/refseq/viral/assembly_summary.txt

ftp://ftp.ncbi.nlm.nih.gov/genomes/refseq/archaea/assembly_summary.txt

Downloading “fna” file using “ftp_path” item of each assembly accession line. For example:

ftp://ftp.ncbi.nlm.nih.gov/genomes/all/GCF/000/189/935/GCF_000189935.1_ASM18993v2/GCF_000189935.1_ASM18993v2_genomic.fna.gz

All references were downloaded using an in-house script:

https://github.com/hitbc/deSAMBA-meta/blob/master/download

We used this script to download the reference sequences via the following command lines:

```
bash ./download -P 10 -o $DOWNLOAD -d bacteria refseq
bash ./download -P 10 -o $DOWNLOAD -d viral refseq
bash ./download -P 10 -o $DOWNLOAD -d archaea refseq
```

All reference files were combined before the construction to the index:

```
find $DOWNLOAD/ -name "*.fna"     | xargs -n 1 cat > ref.fa
```

**2.2 Read datasets and ground truth**

We downloaded the real sequencing datasets from NCBI Sequence Read Archive (SRA). 145 datasets from various bacteria, viral or archaea were employed. It is worthnoting that for all the datasets, only the reads longer than 1000 bp were used for the benchmark.

The ground truth of a dataset was got from NCBI SRA database, and the true organism of a dataset is shown in the "Organism" label:

https://trace.ncbi.nlm.nih.gov/Traces/sra/?run= [ SRA accession]

The taxonomy and genus labels of a dataset are got from NCBI taxonomy database:

https://www.ncbi.nlm.nih.gov/Taxonomy/Browser/wwwtax.cgi

**2.3 Command lines used for read classification**

1) Benchmark for deSAMBA(version 1.0)

Building index:

```
bash ./build-index ref.fa INDEX_DIR
```

Read classification:

```
deSAMBA classify -t 8 -s 62 INDEX_DIR read.fastq > classify.sam
```

2) Benchmark for minimap2 (version 2.16 r922):

Building index for PACBIO data:

```
minimap2 -x map-pb -d ref_pb.mmi ref.fa
```

Building index for NANOPORE data:

```
minimap2 -x map-ont -d ref_np.mmi ref.fa
```

Read classification for PACBIO data:

```
minimap2 -a ref_pb.mmi read.fastq -t 8 > classify.sam
```

Read classification for NANOPORE data:

```
minimap2 -a ref_np.mmi read.fastq -t 8 > classify.sam
```

3) Benchmark for Centrifuge (version 1.0.4)

Building index:

```
centrifuge-build -p 16 --conversion-table ref.map --taxonomy-tree nodes.dmp --name-table names.dmp ref.fa INDEX_DIR
```

Read classification:

```
centrifuge-class INDEX_DIR --out-fmt sam -t -p 8 -U read.fastq > classify.cen
```

4) Benchmark for KAIJU (version 1.6.3)

Building index:

```
kaiju-makedb -s refseq
```

Read classification in MEM mode:

```
kaiju -t node.dmp -f kaiju_db_refseq.fmi -v -a mem -i read.fastq -z 8 > classify.kai
```
